# FunOrder 2.0 – a fully automated method for the identification of co-evolved genes

**DOI:** 10.1101/2022.01.10.475597

**Authors:** Gabriel A. Vignolle, Robert L. Mach, Astrid R. Mach-Aigner, Christian Derntl

## Abstract

Coevolution is an important biological process that shapes interacting species or even proteins – may it be physically interacting proteins or consecutive enzymes in a metabolic pathway. The detection of co-evolved proteins will contribute to a better understanding of biological systems. Previously, we developed a semi-automated method, termed FunOrder, for the detection of co-evolved genes from an input gene or protein set. We demonstrated the usability and applicability of FunOrder by identifying essential genes in biosynthetic gene clusters from different ascomycetes. A major drawback of this original method was the need for a manual assessment, which may create a user bias and prevents a high-throughput application. Here we present a fully automated version of this method termed FunOrder 2.0. To fully automatize the method, we used several mathematical indices to determine the optimal number of clusters in the FunOrder output, and a subsequent k-means clustering based on the first three principal components of a principal component analysis of the FunOrder output. Further, we replaced the BLAST with the DIAMOND tool, which enhanced speed and allows the future integration of larger proteome databases. The introduced changes slightly decreased the sensitivity of this method, which is outweighed by enhanced overall speed and specificity. Additionally, the changes lay the foundation for future high-throughput applications of FunOrder 2.0 in different phyla to solve different biological problems.

**AUTHOR SUMMARY:** Coevolution is a process which arises between different species or even different proteins that interact with each other. Any change occurring in one partner must be met by a corresponding change in the other partner to maintain the interaction throughout evolution. These interactions may occur in symbiotic relationships or between rivaling species. Within an organism, consecutive enzymes of metabolic pathways are also subjected to coevolution. We developed a fully automated method, FunOrder 2.0, for the detection of co-evolved proteins, which will contribute to a better understanding of protein interactions within an organism. We demonstrate that this method can be used to identify essential genes of the secondary metabolism of fungi, but FunOrder 2.0 may also be used to detect pathogenicity factors or remains of horizontal gene transfer next to many other biological systems that were shaped by coevolution.

## INTRODUCTION

Every form of life known to humankind is subjected to evolution. This process shapes and forms all biological systems on macroscopic and molecular level. Thus, understanding and detecting evolutionary processes substantially contributes to understanding life forms and life itself. An important evolutionary process is the so-called coevolution. This is defined as a “process of reciprocal evolutionary change that occurs between pairs of species or among groups of species as they interact with one another” (1). This definition can be extended to interacting proteins (2), may it be physical interactions or may it be consecutive actions in a metabolic pathway. In this regard, coevolution describes a similar evolutionary process with a similar evolutionary history among interacting proteins and the corresponding genes. In a previous study, we described a semi-automated method for the identification of coevolutionary linked genes, named FunOrder (3). Therein, the protein sequences of an input set of proteins are blasted against an empirically optimized proteome database. The Top 20 results of each search are then compared in a multisequence alignment and a phylogenetic tree is calculated for each input protein. Next, the phylogenetic trees of all proteins are compared pairwise using the treeKO tool. This tool calculates how similar two trees are, and in thus how similar the evolutionary history of two proteins is. The treeKO tool calculates two distances, the strict distance and the speciation distance. Notably, the strict distance had previously been suggested to be more suitable for the detection of coevolution in protein families than the speciation (or evolutionary) distance (4). However, we combined the two distance values to a third measure, the combined distance, in order to consider also the speciation history in the FunOrder method. The strict and the combined distances of all pairwise comparisons were then compiled in two matrices and visualized as heatmaps, dendrograms and two principal component analyses (PCA) were performed. In the final step of this method, the user needed to assess these different visualizations of the underlying data to detect co-evolved proteins (Fig 1A). Please also refer to the original study for a detailed description of this method (3).

**Fig 1.**
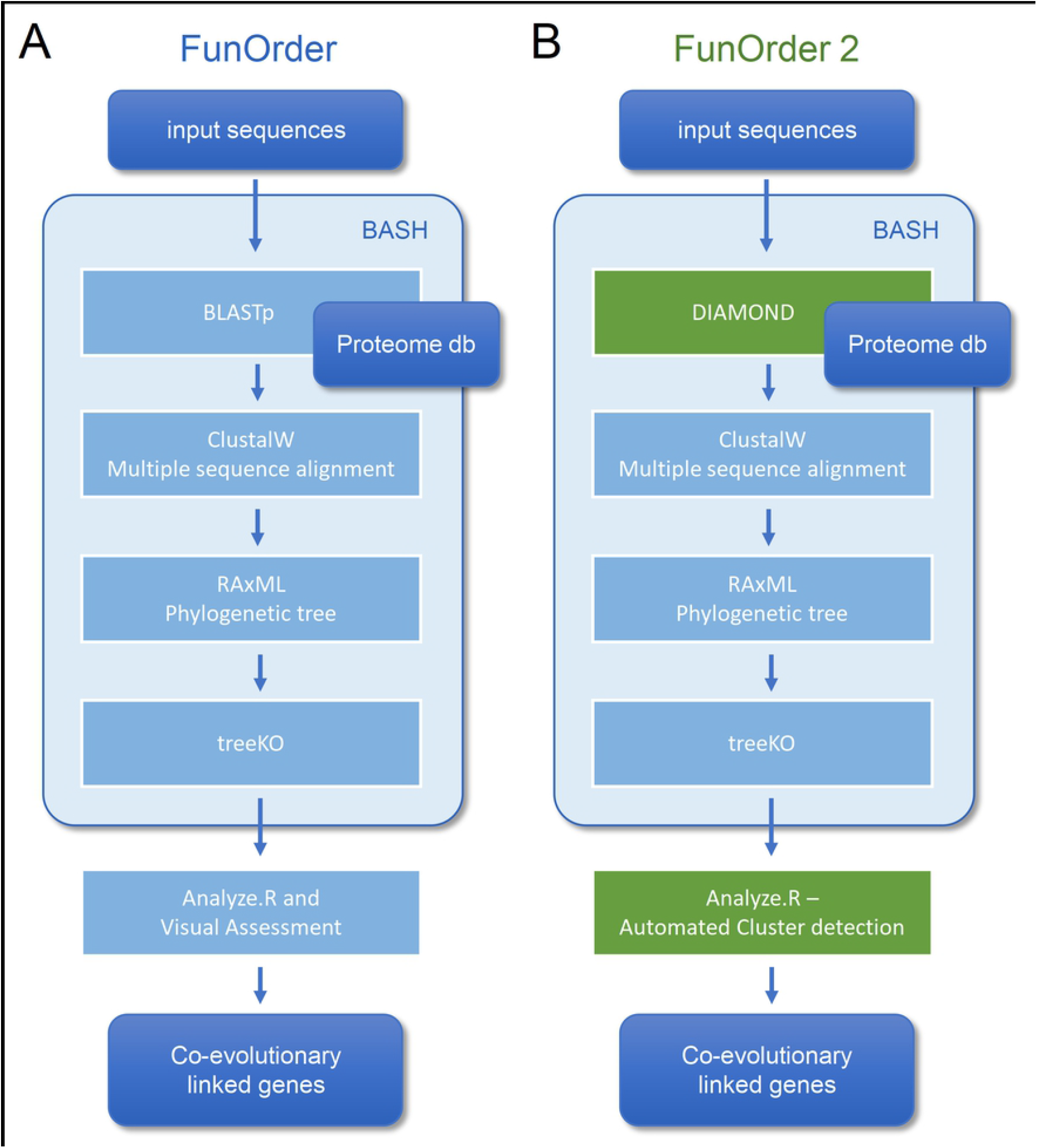
Comparison of the workflow of the original FunOrder method (A) and FunOrder 2.0 (B).

Previously, we demonstrated the functionality and applicability of this method by identifying essential genes in biosynthetic gene clusters (BGCs) of ascomycetes (3). Fungal BGCs contain genes whose corresponding enzymes catalyze the biosynthesis of secondary metabolites (SMs) (5). SMs are a vast group of compounds with different structures and properties that are not necessary for the normal growth of an organism but can be beneficial under certain conditions (6). Notably, many SMs also have medicinal or other useful purposes, such as dyes, food additives, and as monomers for novel plastics (7). However, we can classify the genes in a BGC into biosynthetic genes, further essential genes, and gap genes. The biosynthetic genes encode for enzymes that are directly involved in the biosynthesis of the SM, while the further essential genes encode for transporters (8), transcription factors (9), or resistance genes (10). In contrast, gap genes are not involved in the biosynthesis of the SM despite being co-localized in the BGC (11). Both, the biosynthetic genes and the further essential genes are necessary for the biosynthesis of a SM in the native organisms (12). We could use FunOrder to detect theses essential genes, because they share a similar evolutionary background in many fungal BGCs (3). The FunOrder method contributes to a better understanding of fungal BGCs by adding an additional layer of information. This will support users in the decision which genes should be considered for detailed studies in the laboratory. Importantly, the application of FunOrder is not limited to BGCs from ascomycetes but may be useful to answer any biological problem related to molecular coevolution of genes or proteins. Notably, this requires the compilation and evaluation of a suitable proteome database.

The obvious major shortcoming of the original FunOrder method is the final manual assessment which prevents full automation and high-throughput analyses. Further, the very sensitive but slow BLAST algorithm (13) limits the size of the proteome database used in this method. If this method shall be used for the analysis of coevolution in plants or mammals, larger databases will be needed. In this study we describe an improved version of the method, termed FunOrder 2.0 which overcomes the two mentioned limitations. For an automated detection of co-evolving genes, we determine the optimal number of gene groups in the FunOrder output and then use k-means clustering based on the first three principal components of a PCA. Further, we replace BLAST with the recently published and upgraded DIAMOND tool (14) to enable searching of larger databases and lay the foundation for future different applications of FunOrder 2.0.

## RESULTS

### Integration of the DIAMOND algorithm

The first major improvement of the FunOrder method was the integration of the DIAMOND algorithm (14, 15) for searching the proteome database instead of the previously used BLAST algorithm (13) (Fig 1). This change will allow the usage of larger databases in FunOrder 2.0, since DIAMOND is as sensitive as BLAST, but is faster and is adapted to larger databases (14). With DIAMOND the run time of the first step in the FunOrder pipeline was reduced significantly. For instance, the database search for the lovastatin BGC of *Aspergillus terreus* (lov) (16) took 1 m 25 sec using the original FunOrder method, and 45 sec real time using FunOrder 2.0. This difference will of course be more pronounced and obvious when a larger database is used.

To test, whether the integration of DIAMOND might have altered the ability of FunOrder to detect coevolution, we analyzed the same control gene clusters (GCs) we had previously used to evaluate the original FunOrder method (3) and calculated the internal coevolution quotient (ICQ). The ICQ expresses how many genes in a gene cluster are detected as coevolutionary linked and is calculated subsequently to the treeKO comparison (Fig. 1). Since no other changes have been introduced until this point in the workflow, the ICQ values are a feasible way to compare BLAST and the DIAMOND software. We found only marginal differences between the original FunOrder method (using BLAST) and FunOrder 2.0 (using DIAMOND) (Table S1). For visualization, we compared the ICQ results in a kernel density plot (Fig 2). Therein, the curve for the ICQs of the positive control GCs (BioPath in Fig 2) slightly shifted to the left (higher internal coevolution) compared to the original method, while the curve for the negative control GCs (random GCs in Fig 2) slightly shifted to the right (lower internal coevolution). These results indicate that DIAMOND might be better suited than BLAST within the FunOrder method, as the usage of DIAMOND resulted in a better distinction of the positive and negative control GCs. The curve for the sequential GCs was flattened and broadened compared to the original curve (Fig 2), which can also be explained by the assumed better performance of DIAMOND in this workflow. As the sequential GCs are random loci from different ascomycetes (3), they contain random numbers of co-evolved and independently evolved genes. Consequently, the usage of DIAMOND lowers the ICQ for GCs containing many co-evolved genes and raises the ICQ for GCs with many independently evolved genes compared to the original FunOrder method. This results in the detection of simultaneously more and less coevolution in all sequential GCs and therefore a flattening of the curve in Fig 2. For the benchmark BGCs, we could not observe a drastic change of the height or position of the curve, but a change of the shape with no significant differences of the variance and the mean (File S1). However, the changes of the curves of the random GCs and the BGCs, resulted in a new point of intersection (0.708), which should be considered in the final assessment of fungal BGCs, as described in our previous publication (3).

**Fig 2.**
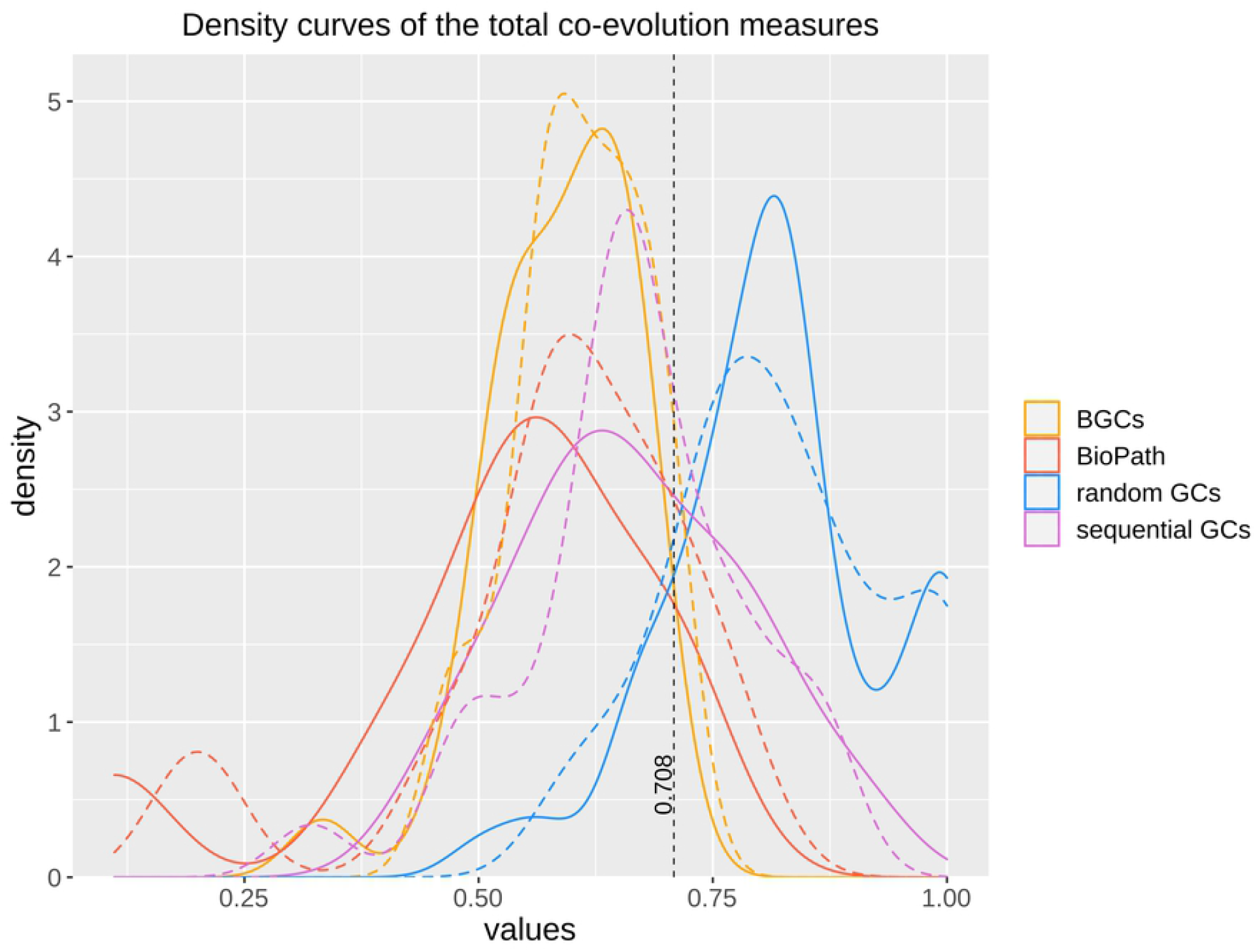
Kernel density plot of the ICQ values for co-evolutionary linked enzymes of different control sets comparing the original FunOrder method (dashed lines) and FunOrder 2.0 (solid lines). BGCs, previously empirically characterized fungal BGCs; BioPath, protein sets of conserved biosynthetic pathways of the primary metabolism; random GCs, randomly assembled protein sets from 134 fungal proteomes; sequential GCs, co-localized genes from random loci of different ascomycetes.

### Automated Cluster definition

As mentioned, a major limitation of the original FunOrder method was the need for a manual assessment of the output, during which the proteins are grouped into clusters based on different data visualizations (Fig 1A). Please refer to our previous method for a detailed description of the procedure (3). To solve this problem, we integrated two R scripts for automatic definition of co-evolved protein groups (or clusters) (Fig 1B). The two R scripts use the first three principal components of the PCA of the strict and the combined distance matrices as input (Fig 1B) and group the proteins by k-means clustering. In the original FunOrder method, only the first two components were considered.

The first R-script for automated protein (or gene) clustering initially determines the optimal number of gene clusters within the first three principal components of the PCAs using the R Package NbClust (17). This package uses different indices and varies the number of clusters, distance measures and clustering methods to determine the optimal number of clusters in a data set based on the majority rule. If the prediction of the optimal number of clusters fails, the second (a back-up) script with a predefined number of clusters is called. The prediction of the optimal number of clusters might fail for instance if the majority rule cannot be applied. As we aim to distinguish biosynthetic, further essential, and gap genes in fungal BGCS, we predefined the number of clusters to 3. Regardless of the script used, the final output is an excel file (Table S2) and a color-coded visualization of the PCA (File S2).

To test how this automated cluster definition compares to the previously performed manual cluster definition, we analyzed the same 30 BGCs as in our previous study. To observe only the influence of the automated cluster definition, we kept the BLAST tool for the initial database search still in place (Fig. 1). Then, we compared the obtained results to those of the previously performed manual analyses (3) (Table 1 and Table S3). In only 5 out of the tested 30 BGCs, the exact same results were obtained (Table S3). In 15 BGCs, the automated cluster definition missed at least one biosynthetic or further essential gene in comparison to the manual assessment, but it could detect more of these essential genes in 5 BGCs. Regarding the gap and extra genes, the automated cluster definition returned less false positives than the manual assessment in 12 BGCs but found more in 4 BGCs. In summary, the automated cluster detection appeared to be more stringent than the manual assessment method, which led to slightly reduced sensitivity but enhanced selectivity (see Table S3 for a detailed statistical analysis).

**Table 1.** **Number of BGCs in which the automated cluster detection (in combination with BLAST or DIAMOND) delivered the same, better, worse, or different results for the given gene categories compared to the manual method.**

|  | BLAST / DIAMOND |  |  |  |
| --- | --- | --- | --- | --- |
|  | same | better | worse | different genes |
| biosynthetic genes | 16 / 17 | 4 / 4 | 10 / 9 | - / - |
| further essential genes | 11 / 11 | 3 / 3 | 5 / 5 | - / - |
| gap genes | 13 / 13 | 8 / 8 | 3 / 3 | 3 / 3 |
| extra genes | 16 / 16 | 5 / 5 | 2 / 3 | 1 / - |

Next, we tested the simultaneous influence of DIAMOND and the automated clustering on the overall performance of FunOrder 2.0 during the analysis of fungal BGCs. To this end, we performed the same comparative analysis of the benchmark BGCs as described above. The results were very similar to the automated analysis using the BLAST analysis (Table 1 and Table S3). In a few cases, the usage of DIAMOND improved the automated cluster definition compared to BLAST, but it remained still more stringent than the manual assessment (Table 1 and Table S3). Fewer biosynthetic genes or further essential genes were detected in 13 of the 30 BGCs by FunOrder 2.0, but also fewer gap or extra genes in 12 BGCs (Table 1 and Table S3). Yet, FunOrder 2.0 clustered more genes together than the original method in other BGCs - to be precise, more essential genes were detected in in 5 BGCs and more gap or extra genes in 4 BGCs compared to the original method (Table S3). The overall enhanced stringency reduced the sensitivity slightly (Table 2) but also improved several statistic measures, including specificity, precision, and the normalized Matthew correlation coefficient (Table 2, in bold). To test if the observed differences have a significant impact on the overall applicability of FunOrder 2.0 in fungal BGCs, we further compared the percentages of correctly identified genes in each BGC between the original FunOrder and FunOrder 2.0 (Tables S3) in an ANOVA (File S1) and found no significant difference. Taken together, we conclude that the introduced changes allow the detection of coevolution between different proteins with an enhanced stringency and precision compared to the original method, and that FunOrder 2.0 can be used to identify essential genes in fungal BGCs.

**Table 2.** Performance comparison of the original FunOrder (3) and FunOrder 2.0 for detecting relevant genes in fungal BGCs. Improved statistical measures are highlighted in bold.

|  | FunOrder<br>essential genes | FunOrder 2.0<br>essential genes | FunOrder<br>biosynthetic genes | FunOrder 2.0<br>biosynthetic genes |
| --- | --- | --- | --- | --- |
| Sensitivity | 0.6349 | 0.6266 | 0.6615 | 0.6564 |
| Specificity | 0.8112 | **0.8541** | 0.8112 | **0.8541** |
| Precision | 0.7766 | **0.8162** | 0.7457 | **0.7901** |
| Negative<br>Predictive Value | 0.6823 | **0.6886** | 0.7412 | **0.7481** |
| False Positive<br>Rate | 0.1888 | **0.1459** | 0.1888 | **0.1459** |
| False Discovery<br>Rate | 0.2234 | **0.1838** | 0.2543 | **0.2099** |
| False Negative<br>Rate | 0.3651 | 0.3734 | 0.3385 | 0.3436 |
| Accuracy | 0.7215 | **0.7384** | 0.7430 | **0.764** |
| F1 Score | 0.6986 | **0.7089** | 0.7011 | **0.7171** |
| Matthews<br>Correlation<br>Coefficient | 0.4524 | **0.4926** | 0.4797 | **0.5242** |
| Normalized Matthews Correlation Coefficient | 0.7262 | **0.7463** | 0.73985 | **0.7621** |
| No-information error rate ni | 0.5084 | 0.5084 | 0.5444 | 0.5444 |

The automation of the cluster detection in FunOrder 2.0 prevents a user bias and improves the overall speed. The analysis of the lovastatin BGC of *Aspergillus terreus* (lov) (16) with 17 genes, took 1 h 19 m 48 sec real time using 22 threads on an Ubuntu Linux system with 128 GB DDR4 RAM with the original FunOrder (excluding manual cluster definition) and 1 h 19 m 58 sec real time with FunOrder2. Notably, the runtime for FunOrder 2.0 includes already the automated detection and grouping of co-evolving genes, which takes an experienced user additional 30 - 45 minutes during the original method.

## MATERIALS AND METHODS

### Changes of the workflow

Within the previously developed work flow (3) (Fig. 1A), we replaced BLAST (13) with DIAMOND (14) for the database search (Fig 1B). The previous and subsequent steps up to the integrated R-scripts remained the same as described in the original FunOrder method (3). Notably, the BLAST algorithm was kept in the software bundle to extract the sequences from the local database and can be used for an optional remote search of the NCBI database (18). The distances measure obtained after the treeKO algorithm were compiled in matrices, which were used as input for three alternative R-scripts (Fig 1B). All R-scripts combine the strict distance matrix and the evolutionary distance matrix to a third distance, the combined distance matrix. These matrices were used for the calculation of the ICQ and as basis for the determination of co-evolved genes. The three scripts differ only in how exactly the co-evolved genes are determined. The first R-script is a revised version of the R-script used in the original FunOrder method (3). It was simplified by removing unnecessary Euclidean distance calculations and the order of the called functions was rearranged in a manner, that all calculations were performed on a single matrix and then on the next. The first matrix to be analyzed is the strict distance matrix followed by the combined distance matrix. Further, we rearranged the order of the visualizations in the output which is saved as “FunOrder_Supplementary_Rplots.pdf”. This output is similar to the original FunOrder method and can be assessed manually as described previously (3).

The second and third R-scripts aim to determine the co-evolved genes automatically. To this end, a PCA is calculated for the strict and the combined distance matrices each. The first three principal components are then considered for defining the clusters by a k-means clustering approach. The difference between the second and the third R-script is the number of clusters used in the k-means clustering approach. In the second R-script, the optimal number of clusters in the first three principal components of the PCAs is determined by NbClust (17) using 28 indices (Table 3). We limited the maximum number of possibly definable clusters to 5 and chose the Ward’s minimum variance method based on the Euclidean distance for optimal cluster search within the NbClust function (19). The third R-script performs a k-means clustering with 3 clusters; it is only called as a back-up if the prediction of an optimal number of clusters in the second script fails. In both cases, the determined clusters are visualized in a color-coded plot of the first two principal components of the PCAs under “FunOrder_clustering_Rplots_pred.pdf” or “FunOrder_clustering_Rplots_defined.pdf” and as table under “cluster_definition_pred.xlsx” or “cluster_definition_3.xlsx”.

**Table 3.** Indices used to determine the optimal number of clusters.

| Index | Reference |
| --- | --- |
| Bale index | (20) |
| Ball index | (21) |
| CCC index | (22) |
| CH index | (23) |
| C-index | (24) |
| DB index | (25) |
| Duda index | (26) |
| Dunn index | (27) |
| Frey index | (28) |
| Friedman index | (29) |
| Gamma index | (30) |
| Gap index | (31) |
| Gplus index | (32, 33) |
| Hartigan index | (34) |
| KL index | (35) |
| Marriot index | (36) |
| McClain index | (37) |
| Pseudot2 index | (26) |
| Ptbiserial index | (32, 38) |
| Ratkowsky index | (39) |
| Rubin index | (29) |
| Scott index | (40) |
| SD index | (41) |
| SDBw index | (42) |
| Silhouette index | (43) |
| Tau index | (32, 33) |
| Tracew index | (44) |
| Trcovw index | (44) |

The software bundle is written in the BASH (Bourn Again Shell) environment and is deposited in the GitHub repository https://github.com/gvignolle/FunOrder (doi:10.5281/zenodo.5118984). Details on all included scripts can be found in the ReadMe file on the GitHub repository. FunOrder 2.0 requires some dependencies, for details and links to all dependencies please refer to the ReadMe file.

### Control gene clusters

For the evaluation of FunOrder 2.0, we used the same control gene clusters (GC) as in the original study (3). As benchmark BGCs, we used 30 previously empirically defined BGCs. As negative controls, we used randomly assembled GCs. As positive control, we used enzymes of conserved metabolic pathways of the primary metabolism. The sequences of all test and control sets are deposited in the GitHub repository https://github.com/gvignolle/FunOrder.

### Calculation of the internal coevolution quotient (ICQ)

“The internal coevolutionary quotient (ICQ) expresses how many genes in a GC or proteins in a protein set are co-evolved according to the previously defined threshold for strict and combined distances within the distance matrices of an analyzed GC (or protein set).”(3) In accordance with the original method the ICQ values were calculated using Equation 1(3).

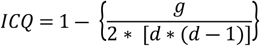

**Equation 1**. ICQ = internal coevolutionary quotient; g = number of strict distances < 0.7 and combined distances <= (0.6 * max value of the combined distance matrix) in all matrices; d = number of genes in the GC (3).

### Performance evaluation

Similar as in the original method we analyzed 30 empirically characterized BGCs to evaluate the ability of FunOrder 2.0 to identify presumably co-evolved essential genes (as defined in Table S3) and to distinguish them from so-called gap genes and genes outside of the defined BGC borders. The genes clustering with the core enzyme(s) were considered as “detected”. As previously described “we counted the total number of (1a) detected essential genes or (1b) detected biosynthetic genes, (2a) not detected essential genes or (2b) not detected biosynthetic genes, (3) detected gap and extra genes, and (4) not detected gap or extra genes in all BGCs, and defined (1a or 1b) as true positives (TP), (2a or 2b) as false negatives (FN), (3) as false positives (FP), and (4) as true negatives (TN)” (3), which were finally used as input for a stringent statistical analysis (3).

## DISCUSSION

The integration of the DIAMOND tool and the automated detection of co-evolved genes improved the run time and total analysis time, allowing high throughput analysis of protein sets and GCs without the risk of a user-bias. In general, the automated cluster definition appears more stringent than the manual assessment, which resulted in improved specificity and precision and a slightly reduced sensitivity during the analysis of fungal BGCs by FunOrder 2.0 compared to the original method. In summary, we consider the integration of a fully automated cluster definition a major improvement, as the advantages (speed, reproducibility, precision) clearly outweigh the slightly reduced sensitivity.

As demonstrated, FunOrder 2.0 can be used to determine the essential genes in fungal BGCs, but this is not the only potential application of FunOrder 2.0. As protein coevolution can be used to predict protein-protein interactions and biosynthetically linked enzymes (3, 45), FunOrder 2.0 may be used to answer many different research questions. Given enough computational time, even complete fungal genomes might be assessed by our method. It is also exciting to speculate if and how FunOrder may be used in other clades of life. A limitation in this regard might be the maximum number of predicted clusters. NbClust limits the number of potential clusters to 15, we further lowered this number to 5 for the analysis of BGCs. This problem might be circumvented by an arbitrary definition of the number of clusters or by consecutive FunOrder analyses, in which a large output cluster is used as input for a new analysis.

FunOrder 2.0 is provided with a database of ascomycete proteomes and can therefore be used for the detection of coevolution of proteins in this fungal division. If other divisions, classes, or even kingdoms shall be analyzed, a suitable new proteome database must be compiled and tested. As mentioned, the integration of the DIAMOND tool enables the integration of larger databases. However, at least 25 different proteomes must be used, because the phylogenetic trees are calculated with a maximum of 20 homologous sequences. Naturally, the proteomes should be of high quality (best RNASeq derived). The proteomes shall be equally distributed among the taxonomic rank to be analyzed but also take the size of the different subordinate ranks into consideration. Put differently, if a division contains 4 small classes, and two large classes, the database should contain proteomes of all six classes, but more from the larger classes than from the smaller classes. The database shall be representative sample of the phylogenetic group to be analyzed. This also means that highly divers phylogenetic groups need to be over-represented in comparison to evolutionary uninventive clades. Further, evolutionary outliers and special clades shall be considered in the database design. For instance, if a phylum contains a family that is the only member of its class, the user needs to decide whether that family shall be part of the proteome database at all, depending on the size and importance of the family. If the family shall be considered, several proteomes need to be included in the database, otherwise the evolutionary distances of the tested proteins might be too large to be successfully evaluated by FunOrder. Any new database must be tested thoroughly according to the procedure we described previously (3). This means, that suitable test gene clusters must be compiled and that meaningful thresholds for the strict and combined distance should be defined. If possible, a test set of target gene clusters should be analyzed and compared to previous results. Please refer to our previous study on how we tested the ascomycete database, determined the thresholds, and tested the applicability of FunOrder for the detection of essential genes in BGCs in ascomycetes (3). A possible short-cut in this procedure might determining the thresholds of strict and combined distance via threshold optimizing (best obtained distinction of positive and negative control gene clusters). Please also refer to the technical guidelines for construction and integration of the database at the GitHub repository https://github.com/gvignolle/FunOrder.

## ACKNOWLEDGEMENT

not applicable

## SUPPORTING INFORMATION

**File S1**. ANOVA for the percentage of correctly detected genes detected by FunOrder and FunOrder 2.0, respectively.

**File S2**. FunOrder 2.0 output of the Lovastatin BGC from *A. terreus* (lov).

**Table S1**. ICQ values of protein sets of conserved metabolic pathways of the primary metabolism (BioPath), sequential GCs and random GCs used in this study.

**Table S2**. FunOrder 2.0 output of the Lovastatin BGC from *A. terreus* (lov).

**Table S3**. Results for the analyses of benchmark BGCs using different versions of the FunOrder method.

## DATA AVAILABILITY

The FunOrder tool, the relevant database, and the sequences and the FunOrder output of the negative control GCs and the positive control BGCs are available in the GitHub repository (https://github.com/gvignolle/FunOrder). We have also used Zenodo to assign a DOI to the repository: 10.5281/zenodo.5118984.

## AUTHOR CONTRIBUTIONS

**GV:** Conceptualization, Data Curation, Formal Analysis, Investigation, Methodology, Software, Validation, Visualization, Writing – Original Draft Preparation

**RM:** Resources, Writing – Review & Editing

**AM:** Resources, Writing – Review & Editing

**CD:** Conceptualization, Funding Acquisition, Methodology, Validation, Project Administration, Supervision, Visualization, Writing – Original Draft Preparation

